# Inferring Multi-Stage Pathway Progression Models from Tumor Phylogenies

**DOI:** 10.64898/2026.05.25.727519

**Authors:** Mikaela Cankosyan, Samin Rahman Khan, Palash Sashittal

**Author notes:** Corresponding authors: Palash Sashittal.

## Abstract

Cancer progression is an evolutionary process driven by the accumulation and selection of somatic mutations, giving rise to genetically diverse subclonal populations within tumors. Understanding the dependencies among mutations and identifying recurrent evolutionary trajectories is critical for understanding cancer progression and informing therapeutic strategies. Recent advances in genomic sequencing and phylogenetic reconstruction now enable large-scale inference of tumor phylogenies, providing detailed representations of intratumor evolutionary histories across patient cohorts. However, modeling cancer progression from these data remains challenging due to extensive inter- and intratumor heterogeneity, often arising from mutations in different genes within the same pathway that confer similar fitness advantages. Existing methods to infer pathway-level progression models summarize each tumor by a single consensus genotype, ignoring intra-tumor heterogeneity, while phylogeny-based methods typically focus on individual mutations and do not model pathways. We introduce PhyloStage, an algorithm for inferring multi-stage pathway-level cancer progression models from large cohorts of tumor phylogenies. PhyloStage represents progression as a partial order over pathways, permitting independent mutations in incomparable pathways while constraining the order of mutations within the same or dependent pathways. The framework also incorporates uncertainty in tumor phylogenies, resolves mutation clusters with unknown ordering, and stratifies patients by progression stage. Applied to a cohort of 120 acute myeloid leukemia (AML) tumor phylogenies, PhyloStage infers progression models that are aligned with known AML progression. On 99 non–small cell lung cancer (NSCLC) patients, PhyloStage stratifies patients into progression stages such that later stages have larger tumor sizes, corroborating phenotypic tumor progression.

## Introduction

Cancer is an evolutionary process in which somatic mutations accumulate and are selected for within a population of proliferating cells. As tumors grow, this process gives rise to genetically distinct subpopulations of cells, a phenomenon known as intratumor heterogeneity, which has been strongly associated with disease progression, therapeutic resistance, relapse, and poor clinical outcomes [32]. Although tumor evolution is inherently stochastic, numerous studies [30, 17, 22] have demonstrated that cancer progression is not arbitrary: across patients and tumor types, recurrent patterns emerge in the frequency and temporal ordering of mutations in genes. These regularities arise because genetic mutations in a cell play a key role in determining its fitness, making certain ordering of mutations more likely than others under selective pressures. Recovering the dependencies that govern these evolutionary dynamics, i.e. inferring cancer progression models, is therefore critical for understanding tumor development and for enabling principled predictions of future evolutionary steps that can inform effective therapeutic strategies [23, 20, 5].

Recent advances in multi-region and single-cell sequencing technologies [8, 36, 21, 42] coupled with computational methods for tumor phylogeny inference [14, 41, 39, 35] have enabled increasingly detailed reconstructions of clonal architecture and evolutionary history within individual tumors. As a result, large cohorts of tumor phylogenies are now available across diverse cancer types, where each phylogeny is represented as a rooted tree with nodes corresponding to cancer clones defined by sets of somatic mutations and edges encoding ancestral relationships between clones. Despite the growing availability of such data, identifying recurrent patterns of evolution across patients remains a central and unresolved challenge.

A key difficulty arises from the extensive mutational heterogeneity observed both within and across tumors. Even among patients with the same cancer type, the specific somatic alterations, including known driver mutations, often differ substantially, weakening the primary signal traditionally used to infer temporal dependencies: the co-occurrence and ordering of mutations across samples. Much of this heterogeneity reflects the fact that cancer evolution is driven by perturbations to signaling, regulatory, and metabolic pathways rather than to individual genes in isolation [38]; consequently, different tumors may acquire functionally equivalent driver events through mutations in distinct genes within the same pathway. Since these pathways and their interactions is only partially characterized [11], there is a pressing need for scalable computational methods that can leverage large cohort-scale phylogenetic data to infer these pathways and their dependencies during cancer progression.

Recent years have seen the development of several computational frameworks for modeling cancer progression from tumor phylogenies across large patient cohorts, which broadly fall into two categories. The first class of methods [12, 2, 31, 16] identify conserved evolutionary trajectories by aggregating ancestral relationships across phylogenies to extract shared partial orders of mutational events, and identifying mutation orderings or evolutionary trajectories that occur across patients more frequently than expected by chance. The second class of methods [24, 13] infers probabilistic cancer progression models that infer the probability of acquiring a mutation given the given the current clonal genotype. These models enable analyses such as identifying mutually exclusive mutations, estimating likely evolutionary trajectories, and predicting future mutations in a tumor. There are two major limitations of the aforementioned methods for inferring cancer progression models from tumor phylogenies. First, these approaches require the input phylogenies to satisfy the *infinite sites assumption* [18], where each mutation arises exactly once in the phylogeny. However, studies have shown evidence of *parallel mutations*, i.e. same mutation occurring multiple times in the phylogeny, across tumors [19]. Moreover, in practice mutations are often annotated at the gene-level. Consequently, several existing methods allow at most one mutation per gene, even though it is common to observe the same gene being independently mutated in multiple subclones along distinct branches of a tumor phylogeny [28]. Second, despite strong evidence that selective pressures in cancer act primarily on pathways rather than single genes [9], neither class of approaches explicitly models cancer progression at the pathway level, limiting their ability to reconcile extensive mutational heterogeneity with conserved evolutionary structure observed across tumor phylogenies.

Existing methods that infer cancer progression models at the pathway level have largely been developed for cross-sectional bulk sequencing data, in which each tumor is summarized by a single binary genotype [33, 4, 27]. These approaches exploit the observation that mutually exclusive mutations often occur in genes belonging to the same functional pathway, reflecting the fact that disruption of a pathway by one mutation typically confers most of the selective advantage and reduces selective pressure for additional mutations in that pathway. Building on this principle, [33] introduced the first pathway-level progression model under a linear evolutionary assumption, which was later generalized to directed acyclic graphs in pathTiMEx [4] using conjunctive Bayesian networks. These methods provided an important conceptual advance by enabling pathway-aware progression modeling and inferring pathway-level dependencies despite extensive gene-level heterogeneity. However, because they rely on a single consensus genotype for each tumor, these approaches fail to account for intratumor heterogeneity. Specifically, mutations that occur in different subclones of the same tumor appear as co-occurring in the consensus genotype, which may lead to incorrect progression models.

To address these limitations, we introduce PhyloStage, a novel framework for inferring pathway-level cancer progression models from large cohorts of tumor phylogenies. We introduce a multi-stage pathway progression model for cancer evolution represented as a *k*-partite partial ordering of pathways. The pathways are partitioned into groups representing distinct stages of cancer progression. Under our model, mutation events are independent if they belong to incomparable pathways, and depend on each other if they belong to the same or comparable pathways under the partial ordering. Our method has several features that are missing in existing methods (see Supp. Table 1 for comparison), such as allowing tumor phylogenies that have parallel mutations (going beyond the restrictive infinite sites assumption), incorporating uncertainty in the tumor phylogenies, resolving mutation clusters with unknown ordering in the phylogenies and stratifying patients based on stages of progression. On real data from 120 trees of acute myeloid leukemia (AML) [28] patients, PhyloStage infers progression models that reveal step-wise progression of that recapitulates known AML biology. PhyloStage also stratifies 99 non–small cell lung cancer (NSCLC) patients [15] in a way that is consistent with phenotypic tumor progression, as patients assigned later stages have larger tumor sizes.

### Multi-stage Pathway Progression Models

We model accumulation of cancer mutations in terms of pathways and dependencies between them that determine the order of occurrence of mutations. Pathways are represented as sets of genes whose products work together in a coordinated manner to perform a specific biological function. Since all genes are essential for the pathway to function, a mutation in only one of the genes is required to disrupt the pathway, which reduces the selective pressure for subsequent mutations in genes in the same pathway. Consequently, tumors tend to harbor at most one driver mutation per pathway, leading to a pattern of mutual exclusivity among genes that participate in the same pathway. Patterns of mutual exclusivity across tumors have previously been used as a signal to identify pathway structure and functional redundancy [4, 37]. In our model as well, we require that mutations in genes belonging to the same pathway are mutually exclusive. Moreover, we assume that pathways are disjoint, so that each gene belongs to at most one pathway.

We introduce a *multi-stage pathway progression model* for cancer evolution, represented by a *k*-partite partial order ≺ on the set 𝒫 of pathways. Specifically, we assume a partition 𝒫 = 𝒫_1_ ∪ 𝒫_2_ ∪ … 𝒫_*k*_ on the set 𝒫 of pathways, where each part 𝒫_*i*_ represents a distinct *stage* of cancer progression. Intuitively, we require a pathway from stage *i* − 1 must be mutated in order for the patient to advance to the next stage *i* of cancer progression. Pathways in the same stage are incomparable, and if pathway *P* precedes pathway *P*′ (i.e. *P* ≺ *P*′) then they must belong to distinct stages *P* ∈ 𝒫_*i*_ and *P*′ ∈ 𝒫_*j*_ such that *i < j*. Under our progression model, a gene in pathway *P* can mutate only if a gene from every pathway *P*′ that precedes *P* (i.e., *P*′ ≺ *P*) has mutated. However, mutations in genes that belong to incomparable pathways can occur independently. While we do not model the probability for occurrence of mutations, our model follows the structural constraints of a conjunctive Bayesian network, which has been used in the past for inferring progression models for cancer and viral evolution [1]. Our model generalizes the previously introduced Pathway Linear Progression Model [33] which corresponds to the special case where each stage contains exactly one pathway, yielding a linear total order on the pathways. The multi-stage pathway progression model can be equivalently represented by a *k*-partite directed acyclic graph *M*_*𝒫*_ corresponding to the Hasse diagram of ≺, where vertices represent pathways 𝒫 and edges represent direct non-transitive precedence relationships (see Fig. 1). We use this representation in the rest of the manuscript.

**Figure 1.**
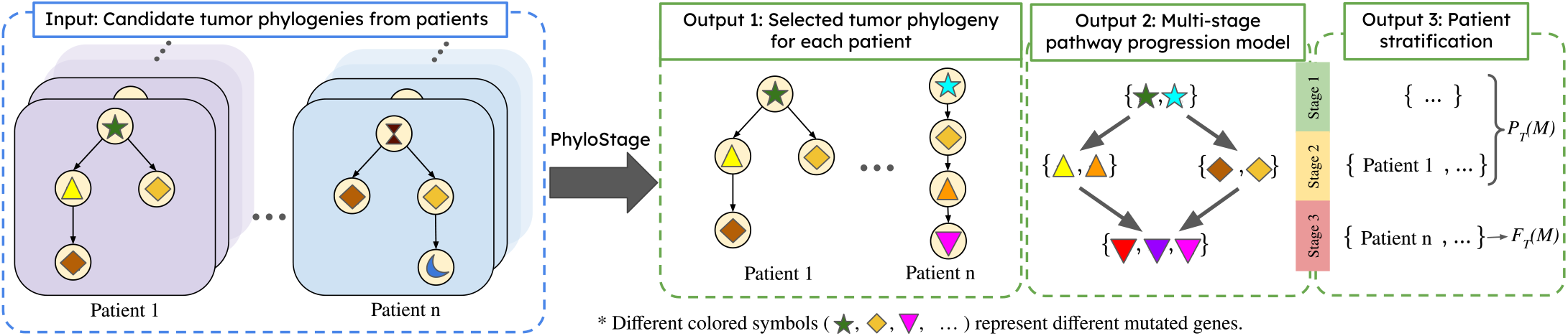
Overview of PhyloStage. PhyloStage takes a family 𝒯 of sets of candidate tumor phylogenies for *n* patients as input. It selects one phylogeny per patient and infers a *k*-stage pathway progression model *M* (*k* = 3 in the example above), that lexicographically maximizes the pair (*F*_𝒯_ (*M*), *P*_𝒯_ (*M*)), where *F*_𝒯_ (*M*) is the number of patients with a tree in which *M* is fully observed and *P*_𝒯_(*M*) is the number of patients with a tree in which *M* is partially observed. Lastly, PhyloStage stratifies the patients by their stage of progression according to the inferred model *M*.

### Inferring Progression Models from Tumor Phylogenies

Suppose we are given tumor phylogenies {*T*_1_, …, *T*_*n*_} of *n* tumors inferred using any of the several existing phylogeny inference methods from bulk or single-cell sequencing data. A tumor phylogeny is a rooted tree, in which vertices represent clones in the tumor and edges represent evolutionary relationship between the clones. Each vertex is labeled by a collection of mutations such that the mutations in a clone are given by the union of all mutations found on the unique path from the root to the clone’s vertex, reflecting the accumulation of mutations over evolutionary time. When a vertex is labeled by multiple mutations, it indicates that the relative temporal ordering of those mutations cannot be resolved from the data, although they are inferred to co-occur in the same clones. Such mutation clusters introduce ambiguity in the evolutionary history and resolving the ordering of mutations within these clusters is a central challenge when identifying conserved evolutionary trajectories across patients. Unlike most existing methods that require the input phylogenies to follow the infinite sites assumption, where each mutation can occur only once, we allow input phylogenies to have parallel mutations, i.e. a mutation can label more than one vertex in the phylogeny. However, we do not allow mutation losses in the phylogenies. If vertex *u*′ lies on the unique path from root to another vertex *u* in *T*, we denote this relationship by *u*′ ≺_*T*_ *u*. If *u* and *u*′ lie on separate branches in *T*, we denote that by *u* ⊥_*T*_ *u*′.

We say that a tumor phylogeny *satisfies* a progression model *M* if it obeys all the mutual exclusivity and ordering constraints induced by *M*. Specifically, the tumor phylogeny must satisfy two conditions. First, the phylogeny *T* must satisfy *pathway exclusivity*, i.e. no two genes *g, g*′ from the same pathway in the progression model can be mutated on the same path from root-to-leaf in *T*. In other words, for any two vertices *u, u*′ with mutation in gene *g* and *g*′ respectively, we must have *u* ⊥_*T*_ *u*′. This ensures that no lineage in *T* may accumulate more than one mutation from the same pathway. Second, *pathway ordering* must be followed, i.e. every vertex *u* with mutation in a gene from pathway *P* that occurs in *T* must be preceded by a vertex *u*′ with mutation in gene *g*′ from every pathway *P*′ such that (*P*′, *P*) is an edge in the progression model *M* (i.e. *u*′ ⪯_*T*_ *u*). We say that a pathway *P* is mutated in phylogeny *T* if at least one gene from *P* is mutated in *T*. Finally, we say that a progression model *M* is *fully observed* in a tumor phylogeny *T* if *T* satisfies *M* and every pathway in *M* is mutated in *T*.

In practice, tumors are sampled at different stages of progression, and thus it is possible that only a subset of the pathways in a progression model are mutated in some of the tumor phylogenies. We formalize this notion through partial observation: a progression model *M* is partially observed in a tumor phylogeny *T* if *T* satisfies *M*, but not all pathways in *M* are mutated in *T*. Thus, while the entire progression model will be observed in tumors at the final stage of progression, partial observation captures tumors that represent intermediate stages of the same underlying progression model.

An important observation is that progression models are not uniquely determined by the set of tumors in which they are observed. In particular, since a pathway is considered mutated in a phylogeny as long as at least one of its genes is mutated, additional genes can be added to a pathway without reducing the number of tumors in which the model is observed. For example, a gene that is never mutated in any of the input phylogenies can be arbitrarily assigned to a pathway without changing the set of phylogenies in which the model is observed.

To eliminate such non-identifiability, we restrict attention to *minimal* progression models. A progression model *M* is *minimal* if the removal of any gene from any pathway strictly decreases the number of tumor phylogenies in which *M* is observed. Intuitively, in a minimal model every gene plays a necessary role in explaining the observed tumors, and no pathway contains superfluous genes whose presence does not affect support for the model.

Due to uncertainty from limited resolution and errors in bulk and single-cell sequencing data, several tumor phylogeny inference methods provide multiple candidate phylogenies that are supported by the observed data for a given patient. Thus, for each patient *i*, we may be given a set 𝒯_*i*_ of plausible tumor phylogenies. Let 𝒯 = {𝒯_1_, …, 𝒯_*n*_} be a family of sets of phylogenies given for *n* patients. For a given progression model *M*, let *F*_*𝒯*_ (*M*) be the number of patients with a tree in which *M* is fully observed and *P*_*𝒯*_ (*M*) be the number of patients with a tree in which *M* is partially observed. When inferring a progression model shared across patients, we must simultaneously (i) select one phylogeny from each set and (ii) identify a progression model that is supported by the selected trees, i.e. maximizes *F*_*𝒯*_ (*M*) and *P*_*𝒯*_ (*M*).

We characterize the complexity of a *k*-stage pathway progression model by three parameters: the number *k* of stages, the number *ℓ* of pathways and the number *d* of dependencies (i.e. the edges in *M*) in the model. Given candidate tumor phylogenies for multiple patients, our goal is to infer a progression model of prescribed complexity that explains as many patients as possible. Since tumors may be sampled at different stages, a model may be fully observed in some patients and only partially observed in others. We therefore adopt a lexicographic objective that prioritizes full observation while using partial observation as a secondary criterion. Intuitively, we seek a model that is fully observed in as many patients as possible (i.e. maximizes *F*_*𝒯*_ (*M*)); among all such models, we prefer one that is also satisfied by (but not necessarily fully observed in) the largest number of additional patients (i.e. subsequently has the highest *P*_*𝒯*_ (*M*)). We formally pose the lexicographical optimization problem as follows:

#### Problem 3.1

(*k*-Stage Pathway Progression Problem) Given a family 𝒯 = {𝒯_1_, …, 𝒯_*n*_} of sets of phylogenies for *n* patients and integers *k, ℓ* and *d*, find a minimal progression model *M* with *k* stages, *ℓ* pathways and *d* dependencies, and select one phylogeny 𝒯_*i*_ ∈ 𝒯_*i*_ for each patient *i*, that lexicographically maximizes the pair (*F*_*𝒯*_ (*M*), *P*_*𝒯*_ (*M*)).

Note that by construction, each stage in a *k*-stage pathway progression model must have at least one pathway and each pathway that is not in the first stage must be dependent on a pathway from the preceding stage. As such, the above problem admits a solution only if *ℓ* ≥ *k* and *d* ≥ *k* − 1. Moreover, the case with *ℓ* = *k* and *d* = *k* − 1 corresponds to a linear progression model [33] in which each stage has exactly one pathway.

An important problem in cancer genomics is to stratify patients based on their stage of cancer progression. We identify the stage of progression of a patient with phylogeny *T* that satisfies the progression model as the highest stage of a pathway that is mutated in *T*. Let *σ*_*M*_ (*T*) denote the stage of progression of phylogeny *T* that satisfies a progression model *M*. If *M* is partially observed in *T*, we have *σ*_*M*_ (*T*) ≥ 1 and *σ*_*M*_ (*T*) = 0 otherwise. Moreover, when a *k*-stage progression model is fully observed in *T*, we have *σ*_*M*_ (*T*) = *k*. Since each pathway belongs to exactly one stage, we formally define the stage of progression for a patient as follows.

#### Definition 1

Suppose a phylogeny *T* of patient satisfies progression model *M* with pathways 𝒫 = {𝒫_1_, …, 𝒫_*k*_}. 𝒯he *stage of progression* of this patient is *σ*_*M*_ (*T*) = max{*i* : there exists *P* ∈ 𝒫_*i*_ mutated in *T* }.

### Characterizing Consistency of Tumor Phylogenies with Progression Models

In this section, we derive a characterization of tumor phylogenies that would satisfy a given progression model *M*. We also show that the *k*-Stage Pathway Progression problem is NP-hard for any number *k* of stages if *k >* 1.

In order for a phylogeny *T* to satisfy a progression model *M*, every clone in *T* must be generated by *M*. We say that a clone is *consistent* with progression model *M* if satisfies two properties: (1) the clone has at most one mutation from each pathway, and (2) if a clone has a mutation from pathway *P*, it must also have mutations from every pathway *P*′ such that (*P*′, *P*) is an edge in *M*. This implies that the pathways mutated in the clone induce a connected subgraph of *M* that includes at least one pathway from stage 1. We derive the following result that provides necessary and sufficient conditions for a phylogeny *T* to satisfy a progression model *M*.

#### Theorem 1

A tumor phylogeny *T* satisfies a progression model *M* if and only if every clone of *T* is consistent with *M*.

The above characterization of tumor phylogenies that satisfy a progression model in terms of their clones allows us to incorporate phylogenies that may have parallel mutations (or genes mutated multiple times in the phylogeny), something that is not possible with most existing methods. Since we do not allow mutation losses in the phylogenies, we can determine the stage of progression of a phylogeny *T* that satisfies a progression model *M*, by examining the pathways mutated in the leaves of *T*.

We show that solving the *k*-Stage Pathway Progression Problem is NP-hard for any value of *k >* 1 even in the restrictive case when the progression model is restricted to be a linear pathway progression model. This corresponds to the case where the number *ℓ* of pathways is *k* and the number *d* of dependencies is *k* − 1.

#### Theorem 2

𝒯he *k*-Stage Pathway Progression Problem is NP-hard for any value of *k >* 1, even when the number *ℓ* of pathways is *k* and the number *d* of dependencies is *k* − 1.

We provide proofs for the theorems in this section in Supp. Sec. 1.

### PhyloStage: An Algorithm to Infer Progression Models and Stratify Patients

In PhyloStage, we leverage the characterization from consistency between tumor phylogenies and progression models derived in the previous section (Theorem 1) to formulate a mixed integer linear program to solve the *k*-Stage Pathway Progression Problem (Prob. 3.1). Given a family 𝒯 of sets of phylogenies for *n* patients and integers *k, ℓ* and *d*, the MILP constructs a *k*-stage pathway progression model *M* with *ℓ* pathways and *d* dependencies that lexicographically maximizes the number *F*_*𝒯*_ (*M*) of patients with a phylogeny in which *M* is fully observed and subsequently the number *P*_*𝒯*_ (*M*) of patients with a phylogeny in which *M* is partially observed. We provides details of the MILP in Supp. Sec. 2.1. Using the progression model *M* inferred by the MILP, PhyloStage stratifies patients based on Definition 1. PhyloStage also performs a permutation-based statistical test to quantify the significance of the inferred progression models (details in Supp. Sec. 2.2).

## Results

### Simulated data

We compare PhyloStage against MASTRO [31], a method that finds statistically significant recurrent gene-level evolutionary trajectories from tumor phylogenies. To our knowledge, no existing method infers pathway-level progression models from cohorts of tumor phylogenies. We evaluate PhyloStage and MASTRO on simulated data generated under a known multi-stage pathway progression model.

We simulate *n* = 120 tumor phylogenies using a 3-stage pathway progression model (i.e. *k* = 3). We base the topology and the number of mutations per clone in our simulated phylogenies on the tumor phylogenies of 120 acute myeloid leukemia patients reported by [28]. Using these fixed tree structures, we simulate mutational assignments via an iterative generative process that places mutations along each phylogeny. The simulation is governed by two parameters. The first parameter, *p*_1_ ∈ [0, 1] denotes the probability that a phylogeny follows the underlying pathway progression model. The second parameter, *p*_2_ ∈ [0, 1] denotes the probability that a mutation is drawn from the progression model (as opposed to being a background mutation), conditioned on the phylogeny following the model. Intuitively, when *p*_1_ = *p*_2_ = 1, the progression model is fully observed in a tumor phylogeny if it contains a clone with at least three mutations (where one mutation will come from each stage), and partially observed otherwise. We vary *p*_1_ ∈ {0.5, 0.8, 1} and *p*_2_ ∈ {0.5, 0.8, 1}, and simulate 20 instances for each combination of parameters. Additional details of the generative procedure are provided in Supp. Sec. 3.

We evaluate the accuracy of PhyloStage and MASTRO for inferring the underlying progression model using two metrics. The first is *gene F1 score*, which measures the harmonic mean of precision and recall (F1 score) of genes in the inferred progression model relative to the ground-truth model used for simulation. The second, *dependency F1 score*, quantifies the harmonic mean of precision and recall of gene–gene dependencies induced by the inferred progression model compared to the ground truth. These results demonstrate that the underlying pathway-level progression structure cannot be reliably recovered by simply aggregating recurrent gene-level trajectories.

PhyloStage outperforms MASTRO across all simulation parameters. For the most challenging simulated instances, with *p*_1_ = *p*_2_ = 0.5, PhyloStage has substantially greater gene F1 score (median 1.0, mean 0.96) and dependency F1 score (median 1.0, mean 0.93) compared to MASTRO (medians 0.67 and 0.35). Similar trends are observed for recall and precision of genes and their dependencies (Supp. Fig. S1). Furthermore, PhyloStage is faster than MASTRO at higher *p*_1_ and *p*_2_ values (median runtime 0.15s vs 0.84s for *p*_1_ = *p*_2_ = 1), while having comparable runtime overall (Supp. Fig. S2). These results demonstrate that the underlying pathway-level progression model cannot be reliably recovered by simply aggregating recurrent gene-level trajectories. Explicitly modeling progression at the pathway level, as done by PhyloStage, enables accurate inference of the underlying progression model.

### PhyloStage infers AML progression models

We use PhyloStage to infer acute myeloid leukemia progression models from *n* = 120 patient tumor phylogenies generated using SCITE [14] in [28]. We focus on linear *k*-stage progression models where number of pathways *ℓ* = *k* and number of dependencies *d* = *k* − 1. Since the median depth of the patient phylogenies is 3, we infer models with *k* = 2 (Supp. Fig. S3) and *k* = 3 stages.

The 3-stage progression model inferred by PhyloStage reveals step-wise progression of AML in 40 patients (Fig. 3, p-value 0.024; significance testing details in Supp. Sec. 5, Supp. Fig. S4a). The first stage is enriched for genes involved in negative regulation of DNA-templated transcription (*p*-value 3.3 × 10^*−*5^, details of pathway enrichment analysis in Supp. Sec. 4). Mutations in this pathway may relieve transcriptional repression, leading to increased or dysregulated RNA production early in disease progression. The second stage is enriched for genes participating in mRNA splicing via the spliceosome (*p*-value 1.1 × 10^*−*5^), suggesting that subsequent alterations disrupt RNA processing and alternative splicing. Together, these first two stages point to a progression trajectory in which transcriptional dysregulation is followed by widespread perturbation of RNA splicing, potentially amplifying transcriptomic instability. The final stage consists of FLT3 and NRAS, key components of a known AML-associated signaling pathway [6], and genes that are also implicated in B-cell differentiation (*p*-value 4.8 × 10^*−*4^). Activating mutations in these genes are well-established drivers of proliferative signaling, consistent with a late-stage transition toward aggressive clonal expansion. Thus, the inferred model suggests a biologically plausible ordering: early disruption of transcriptional control, followed by defects in RNA splicing, culminating in activation of oncogenic signaling pathways. These results illustrate how pathway-level modeling uncovers an interpretable and clinically meaningful progression structure that would be difficult to discern from individual gene-level trajectories alone.

**Figure 2.**
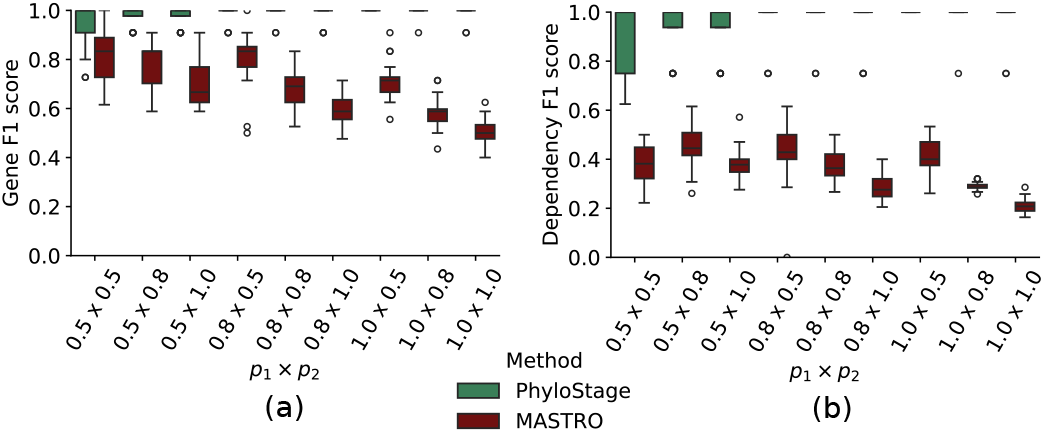
PhyloStage outperforms MASTRO in inferring the underlying progression model from simulated tumor phylogenies. (a) Gene F1 score and (b) dependency F1 score of PhyloStagehy and MASTRO across different values of simulation parameters (*p*_1_, *p*_2_).

**Figure 3.**
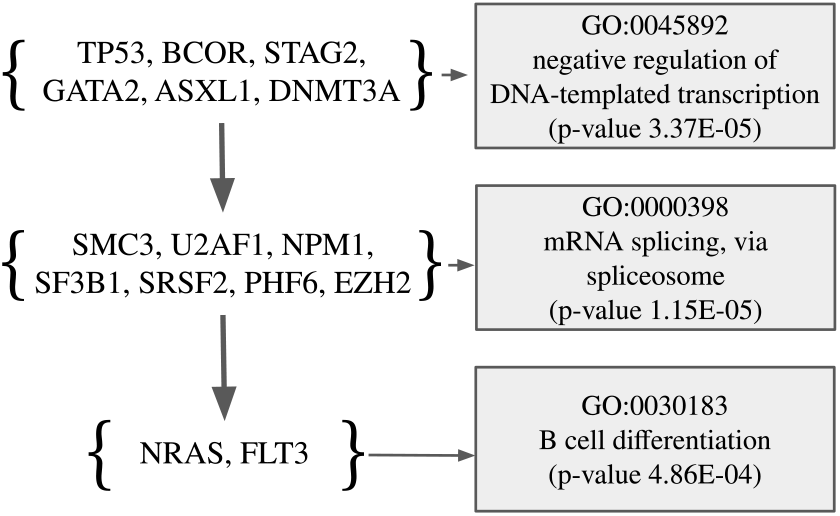
k-stage progression model (*k* = 3) of AML inferred using PhyloStage on 120 patient tumor phylogenies.

### PhyloStage stratifies NSCLC patients by cancer progression stages

We use PhyloStage to infer progression models and stratify 99 non-small cell lung cancer (NSCLC) patients using tumor phylogenies derived from multi-region bulk whole exome sequencing data [15]. Due to uncertainty in the data, this dataset has multiple candidate phylogenies for some patients, with a total of 137 phylogenies across the 99 patients (median 1 tree per patient).

PhyloStage stratifies NSCLC patients into stages of progression in a way that is consistent with observed tumor sizes. We used PhyloStage to infer *k*-stage linear progression models for *k* ∈ {1, 2, 3, 4, 5}. Before running our method, we excluded genes present in fewer than 8 patients to avoid introducing spurious genes into the pathways. We selected the model with *k* = 4 (Fig. 4b, p-value 0.036; details of significance testing in Supp. Sec. 5, Supp. Fig. S4b), since it exhibits a local maximum in total number *F*_*𝒯*_ (*M*) + *P*_*𝒯*_ (*M*) of patients for which the model was partially or fully observed (Fig. 4a). Under this model, 37 patients were stratified into stage 1, 2 into stage 2, 1 into stage 3 and 11 patients into stage 4 (the remaining 48*/*99 patients did not satisfy the inferred progression model). Notably, patients classified as stage 4 exhibited significantly larger tumor sizes than those classified as stage 1 (median 40.0 vs 28.0, *p* = 0.044, Mann-Whitney test [25]). This observation suggests that the progression stages inferred from genotypic data by PhyloStage are consistent with the phenotypic progression of tumors.

**Figure 4.**
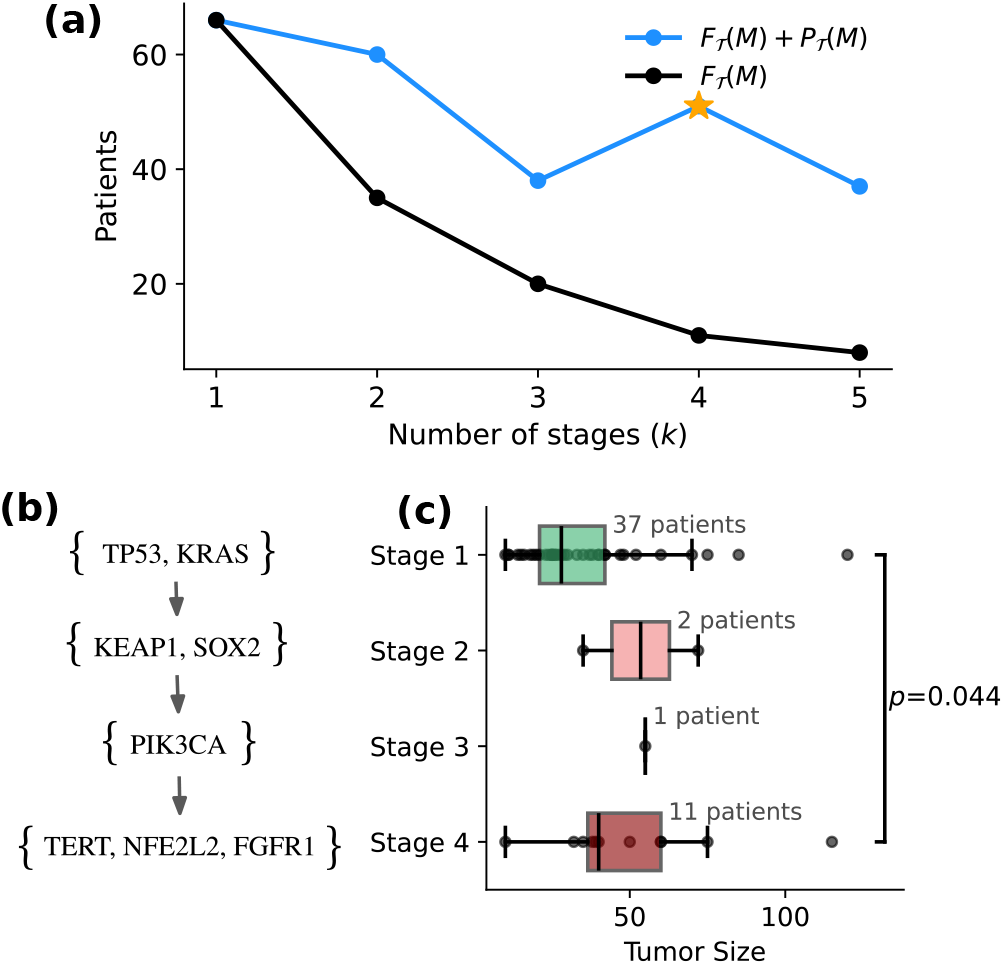
PhyloStage stratifies NSCLC patients into stages consistent with tumor size. (a) Number of patients that satisfy the inferred *k*-stage progression model as a function of *k*; the selected *k* = 4 model is marked with a star. (b) 𝒯he inferred 4-stage progression model. (c) Distribution of tumor sizes for patients assigned to stages 1–4.

## Discussion

In this work, we introduce PhyloStage, novel framework to infer pathway-level cancer progression models from large cohorts of tumor phylogenies and for stratifying patients based on stages of progression. Underpinning our work is a new multi-stage pathway progression model that represents cancer progression as a *k*-partite partial order on a set of pathways. Each stage consists of one or more pathways, and directed dependencies between pathways enforce that at least one pathway in a preceding stage must be altered before progression to a subsequent stage. This formulation provides a biologically interpretable representation of stepwise tumor evolution at the pathway level. We formalize the problem of inferring a *k*-stage pathway progression model that maximizes the number of patients in which the model is fully or partially observed. We show that this problem is NP-hard and formulate a mixed integer linear program to solve it. Through extensive simulations, we demonstrate that PhyloStage scales to large datasets and accurately recovers the underlying pathway-level progression model, even in scenarios where existing approaches fail to identify consistent gene-level evolutionary trajectories. We further apply PhyloStage to tumor phylogenies from 120 acute myeloid leukemia (AML) patients and 99 non-small cell lung cancer (NSCLC) patients. The inferred progression models recapitulate well-established cancer pathways and enable stratification of patients by stage of progression in a manner that is consistent with observed tumor sizes.

There are several avenues for future research. First, our approach depends on the accuracy of input tumor phylogenies. Errors in these phylogenies, such as incorrect placement of mutations, may affect progression model inference. An interesting direction for future work is to infer progression models directly from sequencing data rather than from reconstructed phylogenies. Second, while our framework relaxes the infinite sites assumption and accommodates parallel mutations, it currently does not model mutation losses, which may occur in cancer due to copy-number deletions [29]. Extending PhyloStage to handle both parallel mutations and mutation losses would allow us to leverage phylogenies built under more comprehensive models of cancer evolution [40, 7, 3, 35]. Third, the current model is deterministic and assumes strict mutual exclusivity among mutations within the same pathway. It does not explicitly incorporate probabilistic modeling of mutation acquisition or prediction of future events. Future extensions could relax these assumptions, allowing for probabilistic multi-stage progression models. We envision that as genomic sequencing of cancer tumors becomes increasingly routine in both research and clinical settings, progression-modeling approaches such as PhyloStage will play an important role in elucidating the evolutionary principles underlying cancer development and in bridging genomic progression patterns with clinical decision-making.

## Data availability

Code and data generated in this work are available at https://github.com/sashittal-group/PhyloStage.

## Supplementary Information for

### (1) Proofs

#### Proof of theorem 1

We prove that a tumor phylogeny *T* satisfies a progression model *M* if and only if every clone of *T* satisfies *M*. We restate the theorem here for completeness.

**Theorem**. *A tumor phylogeny T satisfies a progression model M if and only if every clone of T satisfies M*.

*Proof* As stated earlier, a tumor phylogeny *T* satisfies a progression model *M* if and only if it obeys the pathway exclusivity and pathway ordering imposed by *M*. Specifically, there are two conditions: (1a) there are no two genes *g, g*′ from the same pathway *P* in *M* such that *g* and *g*′ are mutated on the same path from root-to-leaf in *T*, and (2a) for all pathways *P, P*′ in *M* such that *P*′ ≺ *P*, for every node *u* in *T* containing a gene *g* in pathway *P*, there must be a node *u*′ containing a gene *g*′ in pathway *P*′ such that *g*′ ≺_*T*_ *g*. For a clone *C* to satisfy a progression model *M*, it must (1b) have at most one mutation from each pathway, and (2b) if a clone contains a mutation in gene *g* from pathway *P*, must also have a mutation from a gene *g*′ in every pathway *P*′ such that *P*′ ≺ *P*.

(⇒) We start by showing that if a tumor phylogeny satisfies a progression model *M*, all its clones satisfy *M*. We prove this by contradiction. Assume that at least one clone *C* in *T* doesn’t satisfy *M*, even though *T* satisfies *M* (i.e. conditions (1a) and (2a) are satisfied). It must be the case that either (1b) or (2b) is violated. If (1b) is violated, *C* has mutations in multiple genes *g, g*′ from the same pathway *P*. However, this would mean that (1a) is violated, because in the path from the root to the node associated with *C*, both *g* and *g*′ are mutated, which is a contradiction. If (2b) is violated, *C* has a mutation in gene *g* from pathway *P*, but not a mutation in gene *g*′ from pathway *P*′ such that *P*′ ≺ *P*. Since *C* contains the union of the sets of mutations in each node on the path from the root to the node associated with *C*, it must be the case that there is no gene *g*′ mutated in *T* such that *g*′ ≺_*T*_ *g*. However, this would mean that (2a) is violated, which is a contradiction. As such, if *T* satisfies *M* it implies that every clone *C* satisfies it.

(⇐) Now, assume that *T* doesn’t satisfy *M*, even though every clone in *T* satisfies *M* (i.e. conditions (1b) and (2b) are satisfied). It must be the case that either (1a) or (2a) is violated. If (1a) is violated, there are two genes *g* and *g*′ from the same pathway *P* in *M* such that *g* and *g*′ are mutated on the same path from root-to-leaf in *T*. Since a clone *C* contains all mutations mutated on the path from the root to its node, there is a clone *C* that contains mutations in *g* and *g*′. However, this would mean that (1b) is violated, which is a contradiction. If (2a) is violated, there is at least one pair of pathways *P, P*′ in *M, P*′ ≺ *P*, such that there is a node *u* in *T* that contains a gene *g* in *P*, but no node *v* in *T* containing a gene *g*′ in *P*′ such that *g*′ ≺_*T*_ *g*. Therefore, it must be the case that the clone *C* corresponding to the node in which a mutation in *g* occurs contains a mutation in *g* but not a mutation in *g*′. However, this would mean that (2b) is violated, which is a contradiction. Therefore, if every clone *C* satisfies *M*, then *T* satisfies *M*. □

#### Complexity Proof

We prove that the *k*-stage pathway progression problem is NP-hard for any value of *k*, even if the number *ℓ* of pathways is *k* and the number of *d* of dependencies is *k* −1. We prove this by reduction from the edge deletion graph bipartization problem to the *k*-stage pathway progression problem. Edge deletion graph bipartization problem, which is a known NP-hard problem [10], is stated as follows: given a graph *G* = (*V, E*), find the smallest set of edges to be removed from *E* such that the resulting graph is bipartite. This is equivalent to finding a bipartition {*V*_1_, *V*_2_} of vertices *V* such that the number of edges *E*_*V*_1 between vertices in *V*_1_ and edges between vertices in *V*_2_ is minimized.

##### Theorem

*The k-stage pathway progression problem is NP-hard for any value of k >* 1, *even if ℓ* = *k and d* = *k* − 1.

*Proof* We prove this by constructing a polynomial-time reduction from the edge deletion graph bipartization problem. Let graph *G* = (*V, E*) with vertices *V* and edges *E* be an instance of the edge deletion graph bipartization problem. We assume that each vertex in *V* has at least one edge (i.e. there is no vertex that has no edge). We construct an instance of *k*-stage pathway progression problem for any value of *k* as follows. For each edge (*u, v*) in *E*, we construct a phylogeny *T*_*u,v*_ that is a linear chain with *k* − 1 vertices, where the first *k* − 2 vertices are labeled by genes *g*_1_, …, *g*_*k−*2_ and the leaf vertex is labeled by {*u, v*}. As such, the number *n* of patients is |*E*|, with one phylogeny per patient, and the number of genes is *k* − 2 + |*V* |. We will show that there exists a solution to the constructed instance of *k*-stage pathway progression problem with number *F*_*𝒯* (*M*)_ of patient phylogenies in which *M* is fully observed equal to *r* if and only if removal of |*E*| − *r* edges can make *G* into a bipartite graph.

Consider progression model *M* with pathways 𝒫 = {*P*_1_, …, *P*_*k*_} as a solution of the *k*-stage pathway progression problem. Since each tree has the same *k* − 2 vertices with one mutation on each, the solution will have *P*_*i*_ = {*g*_*i*_} for *i* ∈ {1, …, *k* − 2}. The remaining pathways we will have mutations corresponding to the vertices in *V*. First, we will show that {*P*_*k−*1_, *P*_*k*_} form a bipartition on *V*. Note that since a gene can belong to at most one pathway, we have *P*_*k−*1_ ∩ *P*_*k*_ = ∅. We need to show that *P*_*k−*1_ ∪ *P*_*k*_ = *V*. We will show this by contradiction. Let there be vertex *v* ∈ *V* that is not in *P*_*k−*1_ ∪ *P*_*k*_ and let *V*_1_ be vertices in *P*_*k−*1_ and *V*_2_ be the vertices in *P*_*k*_. Let *E*_1_ be edges from *v* to vertices in *V*_1_ and *E*_2_ be edges from *v* to vertices in *V*_2_. Note that *M* is not fully observed in any of the phylogenies corresponding to edges *E*_1_ and *E*_2_. However, adding *v* to either *P*_*k−*1_ or *P*_*k*_ will increase the number *F*_*𝒯*_ (*M*) by |*E*_2_| and |*E*_1_|, respectively. This contradicts the claim that *M* is a solution of the *k*-stage pathway progression problem. The bipartition {*V*_1_, *V*_2_} corresponding to the pathways *P*_*k−*1_ and *P*_*k*_ minimize the number of edges *E*_*V*_1 and *E*_*V*_2 and therefore provide a solution to the edge deletion graph bipartization problem.

Consider a solution to the edge deletion graph bipartition {*V*_1_, *V*_2_} such that the number of edges in *E*_*V*_1 and *E*_*V*_2 is minimized, i.e. |*E*_*V*_1 | + |*E*_*V*_2 | is minimized. We construct a progression model with a total ordering of *k* pathways *P*_1_, …, *P*_*k*_, where *P*_*i*_ = {*g*_*i*_} for *i* ∈ {1, …, *k* − 2}, *P*_*k−*1_ = *V*_1_ and *P*_*k*_ = *V*_2_. This progression model is fully observed in phylogenies corresponding to every edge between vertices in *V*_1_ and *V*_2_. It is not partially or fully observed in phylogenies corresponding to edges in *E*_*V*_1 and *E*_*V*_2. Since |*E*_*V*_1 | + |*E*_*V*_2 | is minimized, we have that the number *F*_*𝒯*_ (*M*) of phylogenies in which *M* is fully observed is maximized. □

### (2) PhyloStage: Algorithmic Details

#### (2.1) Mixed Integer Linear Program Formulation

We start by encoding the input data 𝒯 in three binary matrices: patient-tree matrix *D*^*pt*^, the tree-clone membership matrix *D*^*tc*^ and the clone-gene membership matrix 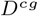. Each entry 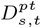 denotes if patient *s* has phylogenetic tree *t*, 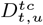 denotes if tree *t* has clone *u* and 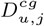 denotes if clone *u* has has a mutation from gene *j*. Let *n*_patients_ be the number of patients, *n*_trees_ be the total number of trees across all patients, and *n*_clones_ be the number of clones across all trees.

We introduce a binary variable *a*_*r,i*_ to denote membership of pathway *i* in stage *r*, and *b*_*i,j*_ for membership of gene *j* in pathway *i*. Since each pathway must have at least one gene and each stage must have at least one pathway, we impose the following constraints

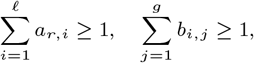

for each stage *r* ∈ [*k*] and each pathway *i* ∈ [*ℓ*]. Moreover, since each gene belongs to at most one pathway, for each gene *j* ∈ [*g*] we impose

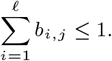

We introduce a binary variable *c*_*u,i*_ to denote mutation of pathway *i* in clone *u*. Since a pathway is mutated in a clone if the clone contains at least one mutation in a gene in the pathway, we impose the following constraints for each clone *u* ∈ [*n*_clones_], each pathway *i* ∈ [*ℓ*], and each gene *j* ∈ [*g*]

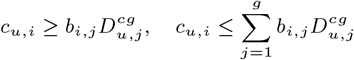

We introduce a binary variable *e*_*i,i*_*′* to denote a dependency of pathway *i′* on pathway *i* (i.e., *i* ≺ *i′*). Since the number of dependencies is given by *d*, we impose

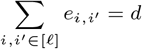

Since a pathway belongs to exactly one stage, for each pathway *i* ∈ [*ℓ*] we impose

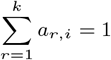

Because each stage needs to occur before the next one, if a pathway has no dependencies, it is in the first stage. So, for each pathway *i* ∈ [*ℓ*] we impose

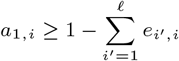

Because dependencies can only be between pathways in consecutive stages, for each stage *r* ∈ [*k* − 1] and each pair of pathways *i, i′* ∈ [*ℓ*] we impose

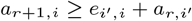

Because there is exactly *k* stages, there cannot be a pathway that is dependent on a pathway in the last stage. So, for each pathway *i* ∈ [*ℓ*] we impose

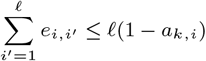

We introduce a binary variable *f*_*u*_ to denote clone *u* satisfying *M*. Because a clone cannot satisfy *M* if it contains multiple mutations that are in the same pathway, for each clone *u* ∈ [*n*_clones_] and each pathway *i* ∈ [*ℓ*] we impose

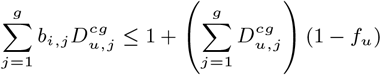

Because a clone cannot satisfy *M* if a pathway *P* is mutated in it but a pathway *P ′* that *P* is dependent on is not mutated, for each clone *u* ∈ [*n*_clones_] and each pair of pathways *i, i′* ∈ [*ℓ*] we impose

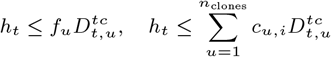

We introduce a binary variable *h*_*t*_ to denote full observation of *M* in tree *t*. For *M* to be fully observed in a tree, all clones present in the tree must satisfy *M*, and all pathways must be mutated in *t*. So, for each tree *t* ∈ [*n*_trees_] and each clone *u* ∈ [*n*_clones_] we impose

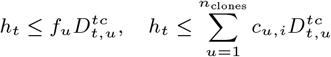

We introduce a binary variable *h′t* to denote partial observation of *M* in tree *t*. For *M* to be partially observed in a tree, all clones present in the tree must satisfy *M*, and at least one pathway must be mutated in the tree. So, for each tree *t* ∈ [*n*_trees_] and each clone *u* ∈ [*n*_clones_] we impose

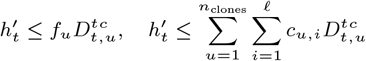

We introduce binary variables *q*_*s*_ and 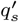 to denote full and partial observation, respectively, of *M* in patient *s. M* is fully or partially observed in a patient if *M* is fully or partially observed in at least one of the patient’s candidate trees. So, for each patient *s* ∈ [*n*_patients_] we impose

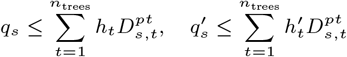

Because we are lexicographically maximizing the number of patients in which *M* is fully observed, followed by the number of patients in which *M* is partially observed, we maximize the following objective function

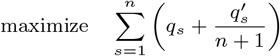

The total number of variables is Θ(*ℓ*(*k* + *g* + *n*_clones_ + *ℓ*)).

#### (2.2) Statistical Significance Testing

To test the statistical significance of our inferred progression models, we start by defining a null distribution of the family 𝒯 = {𝒯_1_, …, 𝒯_*n*_} of sets of phylogenies. We define a null distribution for a phylogeny, and then re-sample each phylogeny under the null distribution. Following the work of [31], we generate a null distribution by permuting the set of mutations for each phylogeny among the nodes of the phylogeny, preserving the topology and the number of mutations in each node. As such, we generate phylogenies assuming that mutations in the tumor are the same but arise independently of any temporal order.

We adapt the procedure used in [34] to estimate the statistical significance of the inferred progression model. We define the score of a progression model as the number of patients in which the progression model is fully observed. We define our test statistic as the the z-score of the score computed on the original dataset against the distribution of scores obtained from datasets drawn from the null distribution. To perform a permutation test, we generate a distribution of z-scores under the null distribution by running PhyloStage on datasets drawn from the null distribution to obtain a progression model, and computing the z-score independently for each one. This null distribution of z-scores represents the extent to which PhyloStage overfits to the noise in the null distribution. Statistical significance of the progression model *M* inferred using the true dataset is quantified by computing the p-value of the z-score for *M*, i.e. the proportion of null z-scores that are greater than or equal to it. Our null hypothesis is that the true dataset was drawn from the null distribution. The p-value for the z-score obtained from the true dataset represents the probability of obtaining a z-score greater than or equal to it under this null hypothesis, so if the p-value is less than a desired significance threshold, then we can reject the null hypothesis, providing evidence that the inferred progression model captures true biological signal.

In summary, we use a permutation test to assess the significance of our progression models, using re-estimation of the test statistic to avoid optimization bias and overfitting. This approach to significance testing is is similar to that used in [34], except that we use z-score of the score as a test statistic rather than the score itself. We do this because the score, i.e. number of consistent patients for a progression model, can be influenced by other factors like number and frequency of mutations in the phylogenies. We chose z-score instead of p-value as the test statistic because for several permuted datasets, we did not have enough samples to resolve very low p-values, causing many of them to be estimated as 0.

### (3) Simulation details

In this section, we describe how we simulate a set of 120 tumor phylogenies using a multi-stage pathway progression model *M*. Each of the 120 simulated tumor phylogenies has a topology corresponding to one of the 120 AML patient tumor phylogenies reported by [28]. We denote the set of mutations present in any of those 120 phylogenies as *A*_*T*_, and the set of mutations present in any of the pathways of *M* as *A*_*M*_. *M* is chosen such that *A*_*M*_ ⊂ *A*_*T*_. All simulations are done with a 3-stage linear pathway progression model: {IDH1, IDH2} → {SFSR1} → {NRAS, KRAS}}. Note that in order to guarantee that every phylogeny can be simulated without error regardless of parameters, it must be the case that |{*A*_*T*_ \ *A*_*M*_ }| ≥ the maximum number of mutations in any of the 120 phylogenies.

Since we are preserving the topology and number of mutations in each node, the process of simulating a phylogeny can be thought of as erasing the identities of all mutations and then sampling them according to the process we will describe. We perform this from top-to-bottom, simulating all nodes at depth *d* before depth *d* + 1. The simulation requires two parameters, *p*_1_ and *p*_2_. Each phylogeny has probability *p*_1_ of “following” *M*. If a phylogeny does not follow *M*, each node samples (without replacement; each mutation can occur at most once in each simulated phylogeny) mutations from *A*_*T*_, preserving the number of mutations in each node.

If a phylogeny follows *M*, since we simulate top-to-bottom, we simulate each stage sequentially. Since *M* is linear, each stage has a single pathway. At a given point in the simulation, we refer to the “current” pathway to mean the latest-stage pathway that has already been mutated in that lineage, and the “next” pathway to mean the pathway in the stage directly following it.

Each node has probability *p*_2_ of advancing stage. If a node does not advance stage, or all pathways in *M* have already been mutated in the phylogeny, or all mutations in the next pathway have already been mutated in the phylogeny (due to branching), then we sample mutations from {*A*_*T*_ \ *A*_*M*_ }. If a node does advance stage, and the other two conditions are not the case, then for each mutation in the node, we sample a mutation from the next pathway. Note that this means that for a node with *x* mutations, those mutations will be sampled from up to *x* consecutive pathways in *M* (we re-check the other two conditions for each mutation since the next pathway changes, so if one of them becomes the case, then it will advance less than *x* pathways). Note that a phylogeny following *M* guarantees that it satisfies *M*, but does not guarantee that it is fully or partially observed.

### (4) Pathway Enrichment Analysis

We used the PANTHER classification system [26] to find the biological processes enriched and significantly overrepresented in the gene sets identified by each pathway in a selected progression model. We used the PANTHER-GO Slim annotation dataset with all genes from the *Homo sapiens* species as reference. The *p*-values of over-representation of each gene set by a biological process were found using Fisher’s exact test.

### (5) Significance Testing on Real Data

We perform the procedure described in Supp. Sec. 2.2 to assess the significance of our results for AML and NSCLC shown in Fig. 3 and Fig. 4. For AML, we obtain a null distribution of z-scores by running PhyloStage with *k* = 3 stages, *ℓ* = 3 pathways, *d* = 2 dependencies, and no restrictions on mutations, on 250 datasets drawn from the null distribution. We estimate the z-score of each progression model by computing a distribution of scores across 1000 null datasets. We then similarly estimate the z-score for the progression model inferred from the true dataset, and compute the p-value of the z-score, which is 0.024 (6/250). The null distribution of z-scores along with the true z-score can be seen in Supp. Fig. S4a. We perform the exact same procedure to assess the significance of our NSCLC progression model, except with different true dataset, tree topologies for null distribution, and settings for Problem 3.1. We use *k* = 4 stages, *ℓ* = 4 pathways, *d* = 3 dependencies, and only consider mutations that are present in at least 8 patients. We obtain a p-value of 0.036 (9/250). The null distribution of z-scores along with the true z-score can be seen in Supp. Fig. S4b.

## (6) Supplementary Figures and Tables

**Figure S1.**
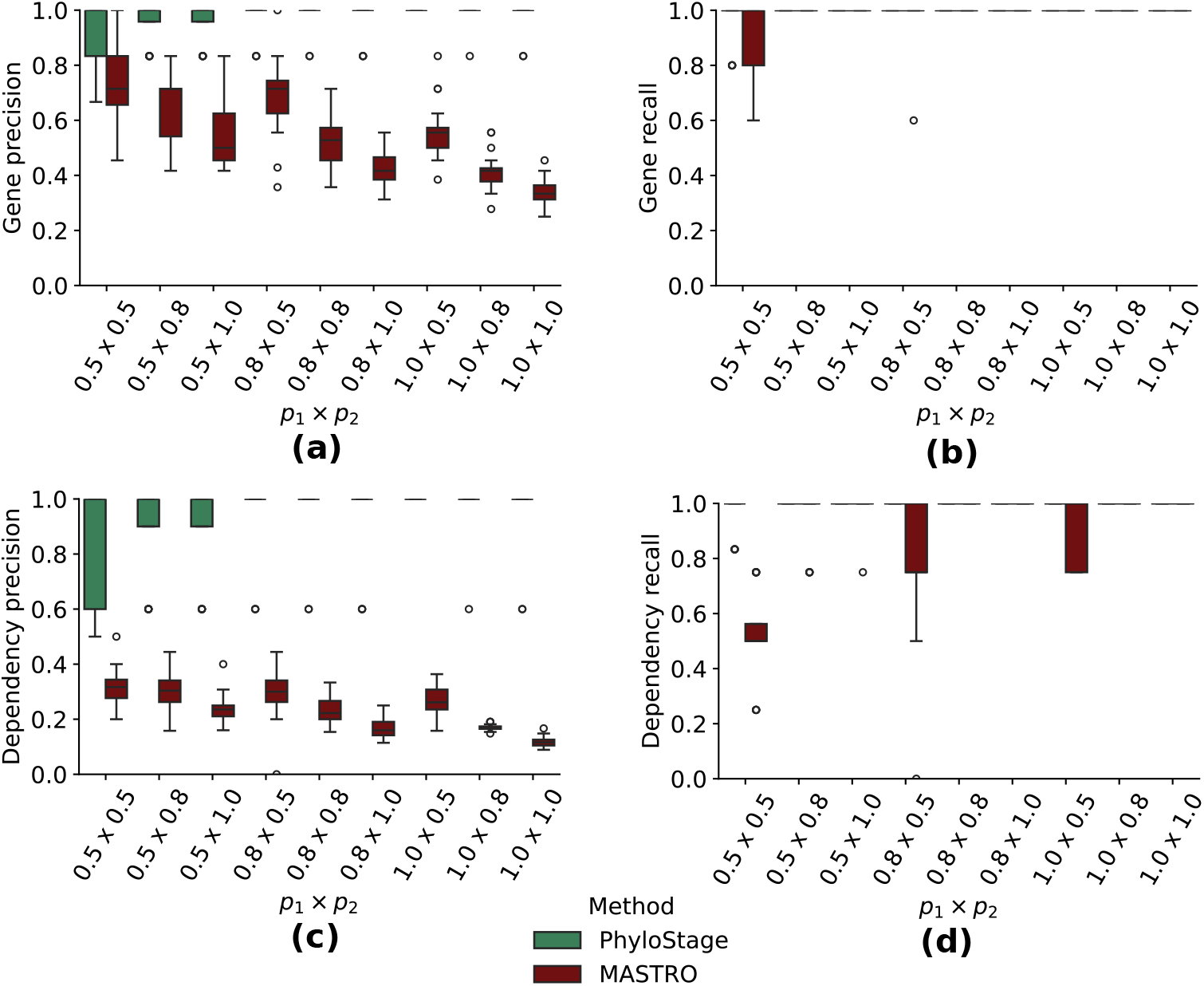
(a) Gene precision, (b) gene recall, (c) dependency precision and (d) dependency recall of PhyloStage and MASTRO across different values of simulation parameters (*p*_1_, *p*_2_).

**Figure S2.**
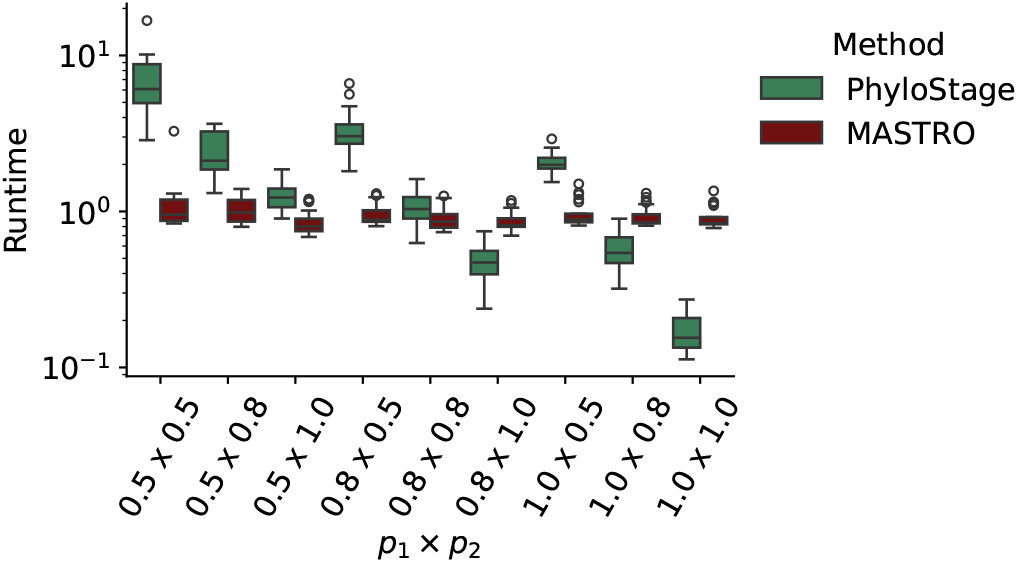
Runtime comparison between PhyloStage and MASTRO across different values of simulation parameters (*p*_1_, *p*_2_).

**Figure S3.**
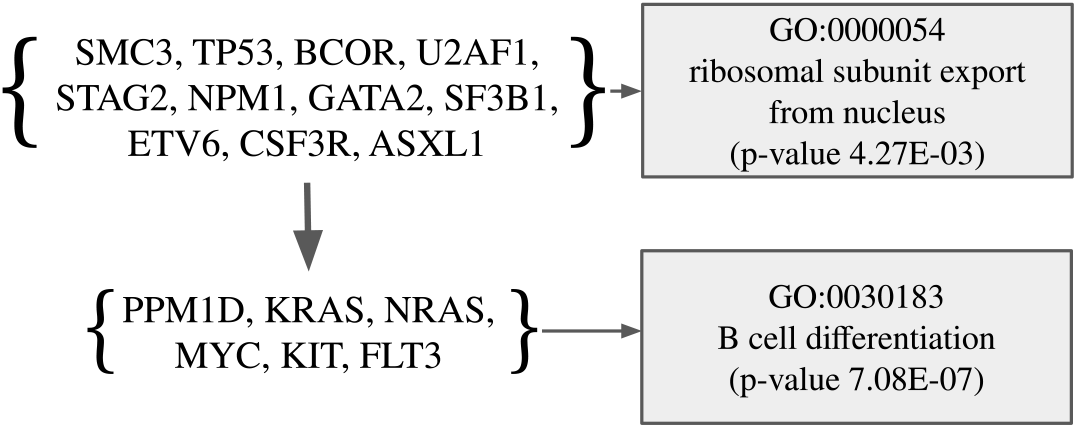
2-stage progression model of AML inferred using PhyloStage on 120 patient tumor phylogenies.

**Figure S4.**
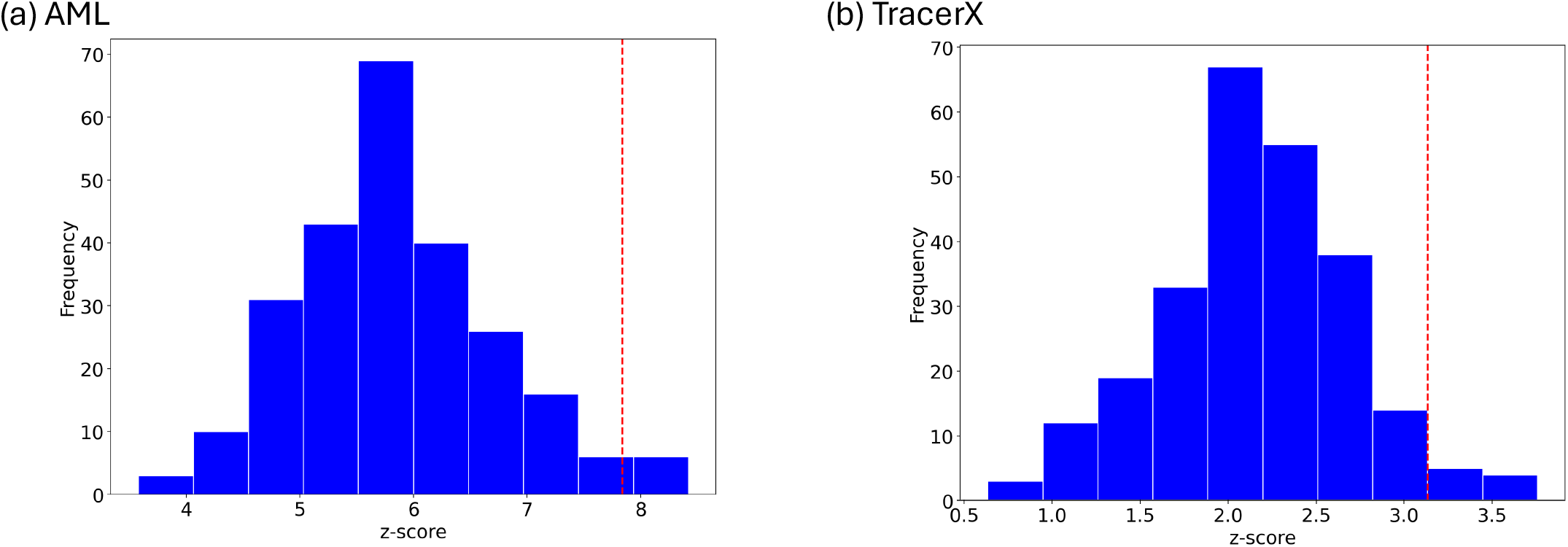
𝒯he distribution of z-scores for the number of patients fully observed in progression models inferred from 250 datasets drawn from the null distribution for (a) AML and (b) 𝒯racerX data. Each z-score is estimated by computing a distribution of number of fully observed patients across 1000 null datasets.

**Table 1.** Properties of existing methods to infer cancer progression model or recurrent trajectories from large cohort of cancer data (either in the form of binary genotype matrix or tumor phylogenies) compared to PhyloStage.

| Method | Pathway | Parallel mutations | Mutation cluster resolution | Tree uncertainty | Patient stratification |
| --- | --- | --- | --- | --- | --- |
| treeMHN [24] | X | X | ✓ | X | X |
| Clomu [13] | X | X | ✓ | ✓ | X |
| RECAP [2] | X | X | ✓ | ✓ | ✓ |
| Conett [12] | X | X | X | ✓ | X |
| MASTRO [31] | X | X | X | X | X |
| POTTR [16] | X | X | ✓ | ✓ | X |
| PathTimeX [4] | ✓ | X | X | X | X |
| PHYLOSTAGE | ✓ | ✓ | ✓ | ✓ | ✓ |

## References

1. Niko Beerenwinkel, Nicholas Eriksson, and Bernd Sturmfels. Conjunctive bayesian networks. Bernoulli, pages 893–909, 2007.

2. Sarah Christensen and Mohammed others. Detecting evolutionary patterns of cancers using consensus trees. Bioinformatics, 36(Supplement 2):i684–i691, 2020.

3. Simone Ciccolella, Mauricio Soto Gomez, Murray D Patterson, Gianluca Della Vedova, Iman Hajirasouliha, and Paola Bonizzoni. gpps: an ilp-based approach for inferring cancer progression with mutation losses from single cell data. BMC bioinformatics, 21(1):1–16, 2020.

4. Simona Cristea, Jack Kuipers, and Niko Beerenwinkel. pathtimex: joint inference of mutually exclusive cancer pathways and their progression dynamics. Journal of Computational Biology, 24(6):603–615, 2017.

5. Ramon Diaz-Uriarte and Claudia Vasallo. Every which way? on predicting tumor evolution using cancer progression models. PLoS computational biology, 15(8):e1007246, 2019.

6. Courtney D DiNardo and Jorge E Cortes. Mutations in aml: prognostic and therapeutic implications. Hematology 2014, the American Society of Hematology Education Program Book, 2016(1):348–355, 2016.

7. Mohammed El-Kebir. SPhyR: tumor phylogeny estimation from single-cell sequencing data under loss and error. Bioinformatics, 34(17):i671–i679, 2018.

8. Charles Gawad et al. Single-cell genome sequencing: current state of the science. Nature Reviews Genetics, 17(3):175–188, 2016.

9. Moritz Gerstung, Nicholas Eriksson, Jimmy Lin, Bert Vogelstein, and Niko Beerenwinkel. The temporal order of genetic and pathway alterations in tumorigenesis. PloS one, 6(11):e27136. 2011.

10. Jiong Guo et al. Compression-based fixed-parameter algorithms for feedback vertex set and edge bipartization. Journal of Computer and System Sciences, 72(8):1386–1396, 2006.

11. Douglas Hanahan. Hallmarks of cancer: new dimensions. Cancer discovery, 12(1):31–46, 2022.

12. Ermin Hodzic, Raunak Shrestha, Salem Malikic, Colin C Collins, Kevin Litchfield, Samra Turajlic, and S Cenk Sahinalp. Identification of conserved evolutionary trajectories in tumors. Bioinformatics, 36(Supplement 1):i427–i435, 2020.

13. Stefan Ivanovic and Mohammed El-Kebir. Modeling and predicting cancer clonal evolution with reinforcement learning. Genome Research, 33(7):1078–1088, 2023.

14. Katharina Jahn, Jack Kuipers, and Niko Beerenwinkel. Tree inference for single-cell data. Genome biology, 17(1):86, 2016.

15. Mariam Jamal-Hanjani et al. Tracking the evolution of non– small-cell lung cancer. New England Journal of Medicine, 376(22):2109–2121, 2017.

16. Sara C Käufler et al. Pottr: Identifying recurrent trajectories in evolutionary and developmental processes using posets. bioRxiv, pages 2026–02, 2026.

17. David G Kent and Anthony R Green. Order matters: the order of somatic mutations influences cancer evolution. Cold Spring Harbor perspectives in medicine, 7(4):a027060, 2017.

18. Motoo Kimura. The number of heterozygous nucleotide sites maintained in a finite population due to steady flux of mutations. Genetics, 61(4):893, 1969.

19. Jack Kuipers, Katharina Jahn, Benjamin J Raphael, and Niko Beerenwinkel. Single-cell sequencing data reveal widespread recurrence and loss of mutational hits in the life histories of tumors. Genome research, 27(11):1885–1894, 2017.

20. Michael Lässig et al. Predicting evolution. Nature ecology & evolution, 1(3):0077, 2017.

21. Devon A Lawson et al. Tumour heterogeneity and metastasis at single-cell resolution. Nature cell biology, 20(12):1349–1360, 2018.

22. Arnold J Levine, Nancy A Jenkins, and Neal G Copeland. The roles of initiating truncal mutations in human cancers: the order of mutations and tumor cell type matters. Cancer cell, 35(1):10–15, 2019.

23. Kamil A Lipinski et al. Cancer evolution and the limits of predictability in precision cancer medicine. Trends in cancer, 2(1):49–63, 2016.

24. Xiang Ge Luo, Jack Kuipers, and Niko Beerenwinkel. Joint inference of exclusivity patterns and recurrent trajectories from tumor mutation trees. Nature communications, 14(1):3676, 2023.

25. Henry B Mann and Donald R Whitney. On a test of whether one of two random variables is stochastically larger than the other. The annals of mathematical statistics, pages 50–60, 1947.

26. Huaiyu Mi and Paul Thomas. Panther pathway: an ontology-based pathway database coupled with data analysis tools. In Protein networks and pathway analysis, pages 123–140.Springer, 2009.

27. Mohammadreza Mohaghegh Neyshabouri and Jens Lagergren. Tomexo: A probabilistic tree-structured model for cancer progression. PLOS Computational Biology, 18(12):e1010732, 2022.

28. Kiyomi Morita et al. Clonal evolution of acute myeloid leukemia revealed by high-throughput single-cell genomics. Nature communications, 11(1):1–17, 2020.

29. Caitlin A Nichols et al. Loss of heterozygosity of essential genes represents a widespread class of potential cancer vulnerabilities. Nature communications, 11(1):2517, 2020.

30. Christina A Ortmann et al. Effect of mutation order on myeloproliferative neoplasms. New England Journal of Medicine, 372(7):601–612, 2015.

31. Leonardo Pellegrina and Fabio Vandin. Discovering significant evolutionary trajectories in cancer phylogenies. Bioinformatics, 38(Supplement 2):ii49–ii55, 2022.

32. Marco Proietto et al. Tumor heterogeneity: preclinical models, emerging technologies, and future applications. Frontiers in Oncology, 13:1164535, 2023.

33. Benjamin J Raphael and Fabio Vandin. Simultaneous inference of cancer pathways and tumor progression from cross-sectional mutation data. Journal of Computational Biology, 22(6):510–527, 2015.

34. Rebecca Sarto Basso, Dorit S Hochbaum, and Fabio Vandin. Efficient algorithms to discover alterations with complementary functional association in cancer. PLoS computational biology, 15(5):e1006802, 2019.

35. Palash Sashittal, Haochen Zhang, Christine A Iacobuzio-Donahue, and Benjamin J Raphael. Condor: tumor phylogeny inference with a copy-number constrained mutation loss model. Genome biology, 24(1):272, 2023.

36. Samra Turajlic et al. Tracking cancer evolution reveals constrained routes to metastases: Tracerx renal. Cell,173(3):581–594, 2018.

37. Fabio Vandin, Eli Upfal, and Benjamin J Raphael. De novo discovery of mutated driver pathways in cancer. Genome research, 22(2):375–385, 2012.

38. Bert Vogelstein et al. Cancer genome landscapes. science, 339(6127):1546–1558, 2013.

39. Jeff A Wintersinger, Stephanie M Dobson, Ethan Kulman, Lincoln D Stein, John E Dick, and Quaid Morris. Reconstructing complex cancer evolutionary histories from multiple bulk dna samples using pairtree. Blood Cancer Discovery, 3(3):208–219, 2022.

40. Hamim Zafar et al. Sifit: inferring tumor trees from single-cell sequencing data under finite-sites models. Genome biology, 18(1):1–20, 2017.

41. Hamim Zafar, Nicholas Navin, Ken Chen, and Luay Nakhleh. Siclonefit: Bayesian inference of population structure, genotype, and phylogeny of tumor clones from single-cell genome sequencing data. Genome research, 29(11):1847–1859, 2019.

42. Haochen Zhang et al. Genomic evolution of pancreatic cancer at single-cell resolution. Nature Genetics, pages 1–11, 2026.

